# The recycling endosome biogenesis machinery coordinates BACE1 endosomal sorting and amyloid-β production

**DOI:** 10.1101/2023.04.05.535676

**Authors:** Jean-Baptiste Brault, Zengzhen Liu, Sabine Bardin, Delphine Ladarré, Vincent Fraisier, Anna Tchenio, Zsolt Lenkei, Jean Salamero, Cédric Delevoye, Bruno Goud, Stéphanie Miserey

## Abstract

Alzheimer’s disease (AD) is the most common form of dementia worldwide. One of AD’s main pathological hallmarks is the cerebral plaque deposits of β-amyloid (Aβ). Aβ is generated through sequential enzymatic cleavage of the amyloid precursor protein (APP). The β-secretase or β-site APP-cleaving enzyme 1 (BACE1) initiates this cleavage and is thus key to regulate Aβ formation. Both APP and BACE1 transit through the endolysosomal system but the exact nature of the compartment(s) where APP cleavage occurs as well as the molecular mechanisms that govern their endosomal sorting remain poorly known. Here we show that RAB11 not only regulates BACE1 transport from early/sorting endosomes (EEs/SEs) and drives the exocytosis of BACE1-containing recycling carriers. Moreover, recycling endosome-associated KIF13A, as well as its closely related homolog KIF13B, which are known RAB11 effectors involved in the biogenesis of recycling endosome (RE) from EEs/SEs, also participate in BACE1 endosomal sorting. Importantly, depletion of KIF13A or KIF13B leads to an increase in Aβ generation. Depletion of the BLOC-1 complex, previously described as an essential partner for KIF13A-dependent RE biogenesis, also induces increased amount of Aβ. Altogether, our findings support a model where the EEs/SEs represent a major organelle for Aβ formation and identify the recycling endosome biogenesis machinery as a master coordinator of BACE1 endosomal sorting and transport.

## Introduction

Alzheimer’s disease (AD) is the most common form of dementia and accounts for 60 to 80 percent of cases worldwide. One of AD’s major pathological hallmark is the cerebral aggregation of amyloid-β peptides (Aβ) in the form of senile plaques. Aβ originates from the sequential cleavage of the transmembrane amyloid precursor protein (APP). The β-secretase or β-site APP-cleaving enzyme 1 (BACE1), a transmembrane aspartyl protease, initiates APP amyloidogenic processing and is thus the rate-limiting enzyme in the process of Aβ generation (Thinakaran and Koo, 2008); (Sinha et al., 1999); (Vassar et al., 1999); (Yan et al., 1999). The amyloid cascade hypothesis posits that Aβ deposition is the root cause of AD. Although this hypothesis is still intensely debated (Herrup, 2015), there is now strong evidence supporting that β-cleavage is essential for the development of the disease (Jonsson et al., 2012).

Both BACE1 and APP are type I transmembrane proteins that traffic from the endoplasmic reticulum along the biosynthetic/secretory pathway (Tan and Gleeson, 2019). After reaching the Golgi complex, newly synthetized BACE1 is likely transported directly to the plasma membrane (PM) (Tan et al., 2020). Previous studies have suggested that APP follows the same route (Haass et al., 2012) but more recent ones indicate that APP is transported from the Golgi complex directly to early endosomes using the AP-4 /Arl5b transport machinery and from there to the PM (Burgos et al., 2010); (Toh et al., 2018); (Tan et al., 2020). Cell surface-associated BACE1 and APP are then internalized and transit through the endolysosomal system (Tan and Gleeson, 2019); (Small et al., 2005).

The convergence of BACE1 and APP at the same subcellular location is critical for the production of Aβ (Das et al., 2013). The exact nature of the compartment(s) where BACE1 cleaves APP remains ill-defined. The prevailing view is that most β-cleavage occurs in early endosomes (EEs) where the acidic pH (pH 4.0-5.0) is optimal for BACE1 enzymatic activity (Shimizu et al., 2008). However, evidence also exists that Aβ is generated predominantly in the TGN (Choy et al., 2012). In addition, the accumulation of APP at the *trans*-Golgi network (TGN), or the redirection of BACE1 to the TGN using a chimeric molecule was shown to increase Aβ production (Chia et al., 2013). From EEs, BACE1 traffics to recycling endosomes (REs) and is recycled back to the PM (Chia et al., 2013); (Udayar et al., 2013); (Buggia-Prévot et al., 2013); (Keable et al., 2022). BACE1 can also be routed to the TGN via the retromer protein complex and GGA proteins, which have been implicated in AD (Small et al., 2005); (Muhammad et al., 2008); (Rogaeva et al., 2007); (Finan et al., 2011); (Wen et al., 2011).

REs are classically characterized by the presence of the RAB11 GTPase (Ullrich et al., 1996; Ren et al., 1998). They consist of an endosomal network of interconnected tubular membrane domains specialized in diverting internalized cargoes towards the plasma membrane. RE tubules originate from vacuolar endosomal sorting domains of EEs or sorting endosomes (SEs) and transport their content along microtubule tracks (Willingham et al., 1984). RE biogenesis requires the coordination of numerous regulators (Goldenring, 2015);(Simonetti and Cullen, 2019). This includes the plus-end-directed kinesin KIF13A which interacts with RAB11 and drives RE tubulogenesis and Tf recycling (Delevoye et al., 2014), as well as RE positioning (spatialization) next to melanosome and cargo transport to melanosome (Delevoye et al., 2009). In addition, BLOC-1 (Biogenesis of Lysosome-related Organelle Complex 1) contributes to KIF13A -dependent tubulation and RE functions (Delevoye et al., 2016);(Jani et al., 2022).

Here we aimed to precise the role of RAB11 and the machinery involved in RE biogenesis in BACE1 trafficking and recycling. We show that RAB11 both controls BACE1 transport at the level of sorting endosomes and participates in the exocytosis of BACE1-containing vesicles. We also provide evidence that KIF13A controls exit of BACE1 from SEs and consequently Aβ production. Altogether, our findings put the RE biogenesis machinery at the center of BACE1 intracellular transport and Aβ production regulation.

## Results and discussion

### BACE1 co-localizes with RAB11A and RAB11B

In order to first unravel the role played by RAB11 in BACE1 intracellular trafficking, Neuro-2a cells were co-transfected with BACE1-GFP and mCherry-RAB11A or mCherry-RAB11B expressing constructs. Of note, these two RAB11 isoforms are expressed in neuronal cells (Goldenring et al., 1994). As previously described (Chia et al., 2013); (Udayar et al., 2013), in control cells co-expressing BACE1-GFP and mCherry, BACE1-GFP was observed in tubulo-vesicular structures spread out all over the cytoplasm (Figure 1A). Consistent with a previously reported function for RAB11 in BACE1 endosomal recycling towards the plasma membrane (Udayar et al., 2013), mCherry-RAB11A and BACE1-GFP co-localized extensively with particular accumulations at the cell periphery (Figure 1A). Similar observations were made upon BACE1-GFP and mCherry-RAB11B co-expression (Figure 1A). The extent of co-localization was quantified using Pearson’s coefficient (Figure S1A-C) and was similar when measured for BACE1-GFP together with mCherry-RAB11A or -RAB11B than when measured for GFP-RAB11A together with mCherry-RAB11A. Co-localization between BACE1-GFP and mCherry-RAB1A (associated with pre-Golgi compartments), used here as a negative control, was lower (Figure S1A-C). Interestingly, when expressed in Neuro2A cells, APP-mCherry was associated with punctate structures not co-localized with GFP-RAB11A (Figure S1B-C) but largely surrounded and/ or closely apposed to LAMP1-GFP positive structures (Figure S1B-D). This strongly suggests that unlike BACE1, APP does not traffic through a RAB11-dependent recycling transport pathway and rather goes to the degradative compartments as previously described (Haass et al., 2012).

**Figure 1:**
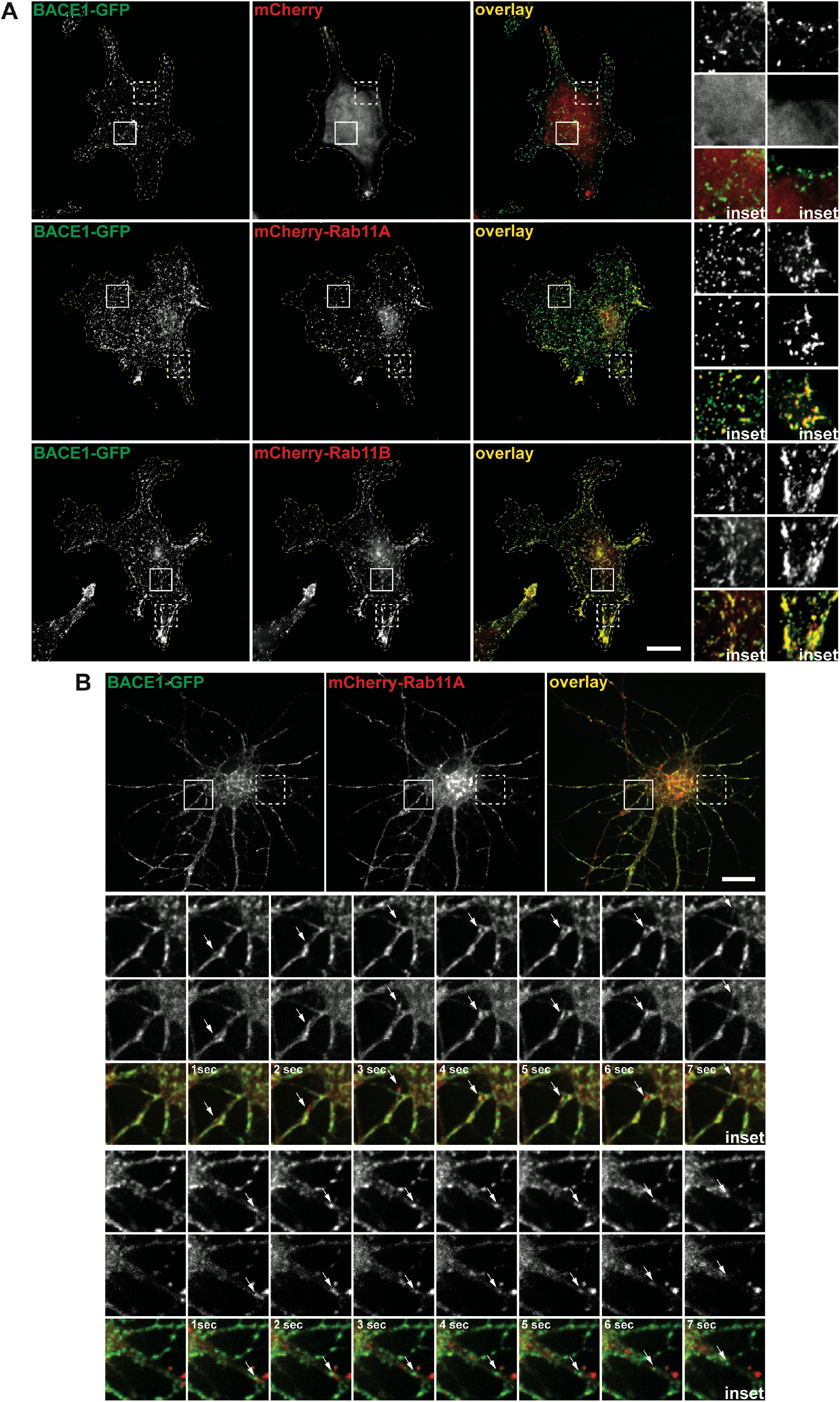
BACE1 co-localizes with RAB11A and RAB11B. **(A)** Neuro-2a cells co-expressing BACE1-GFP and mCherry, mCherry-RAB11A or mCherry-RAB11B. Representative images of cells expressing BACE1-GFP (green) and mCherry, mCherry-RAB11A or mCherry-RAB11B (red) are displayed. Magnified insets from squared boxes are shown on the right. **(B)** Rat hippocampal neurons co-expressing BACE1-GFP and mCherry-RAB11A were imaged every second for 1 min using time-lapse microscopy (see Movie S1). Representative images taken from movies are displayed. Representative images of the dynamic motion of mCherry-RAB11A and BACE1-GFP positive vesicles moving together (arrows). Higher magnification (boxes) of the images taken from time-lapse movies are on the bottom. Bars: 10 µm

To further analyze the dynamic behaviour of BACE1 and RAB11, time-lapse live microscopy was performed in primary rat hippocampal neurons grown at low density and expressing BACE1-GFP and mCherry-RAB11A (Figure 1B). Dynamic BACE1-GFP-containing tubulo-vesicular structures decorated with mCherry-RAB11A could be observed in both the cell soma and in neurites (Figure 1B, Movie S1). Two examples of such events are depicted in still images extracted from movie S1 (Figure 1B) and show the case of transport of a vesicle in dendrites, suggesting that RAB11, in agreement with previous work (Buggia-Prévot et al., 2013) regulates BACE1 vesicular transport in neurons. Altogether these data support a role for RAB11A and RAB11B in BACE1, but not APP, intracellular trafficking.

### RAB11 participates in the exocytosis of BACE1-containing vesicles

RAB11 regulates exocytosis of recycling cargoes such as transferrin (Ullrich et al., 1996); (Takahashi et al., 2012) and langerin (Gidon et al., 2012). Given that RAB11 dynamically associates with BACE1-containing vesicles at the cell periphery, we reasoned that it may regulate BACE1 exocytosis. Therefore, we used a pH sensitive superecliptic pHluorin-tagged BACE1 (pHBACE) construct (Bauereiss et al., 2015) and performed high-resolution Mutli-Angle total internal reflection fluorescence (MA-TIRF) microscopy experiments (Boulanger et al., 2014). HeLa cells co-expressing pHBACE with mCherry-RAB11A or -RAB11B was used to capture concomitant exocytic events, whereas RFP-RAB5A (associated with early endosomes) or mCherry-RAB6A (associated with the Golgi complex) were used as controls. Selected areas (white squares in Figures 2A-B) of still images extracted from movies S2 and S3 obtained by a single TIRF angle are compared with a side view of the corresponding cuboid after 3D reconstruction (Figures 2A-B). The depicted events illustrate the sequence of a RAB11A or a RAB11B positive vesicle docking at the plasma membrane (Figures 2A and 2B, time point 0s628ms) prior to its exocytosis as visualized by the appearance of a bright green signal due to pH sensitivity of pHluorin (Figures 2A, time point 5s130ms and B, time point 37s970ms, arrowheads). Then both red and green signals vanished quickly indicating the termination of the exocytosis (Figures 2A, time point 6s684ms and 9s692ms, and 2B, time point 38s434ms and 40s105ms). We measured the total amount of pHBACE exocytotic events and their co-localization with the different RAB proteins (Figures 2C and 2D). RAB11A or RAB11B co-distributed at the site of pHBACE exocytosis in about 75% of the cases (Figure 2C). On the contrary, RAB5A or RAB6A displayed low co-localization (20% or 8%, respectively), suggesting that RAB11 represents a specific member of the RAB family associated with BACE exocytosis (Figure 2C). Accordingly, RAB11A- or RAB11B-positive total structures associated with pHBACE exocytotic event were over-represented, almost 10-folds higher, relative to RAB5A- or RAB6A-structures (Figure 2D). Altogether, these results show that RAB11, but not other tested RAB proteins, is present at the site of BACE1 exocytosis, strongly suggesting a role in the regulation of the tethering/docking and/or fusion mechanism at the plasma membrane.

**Figure 2:**
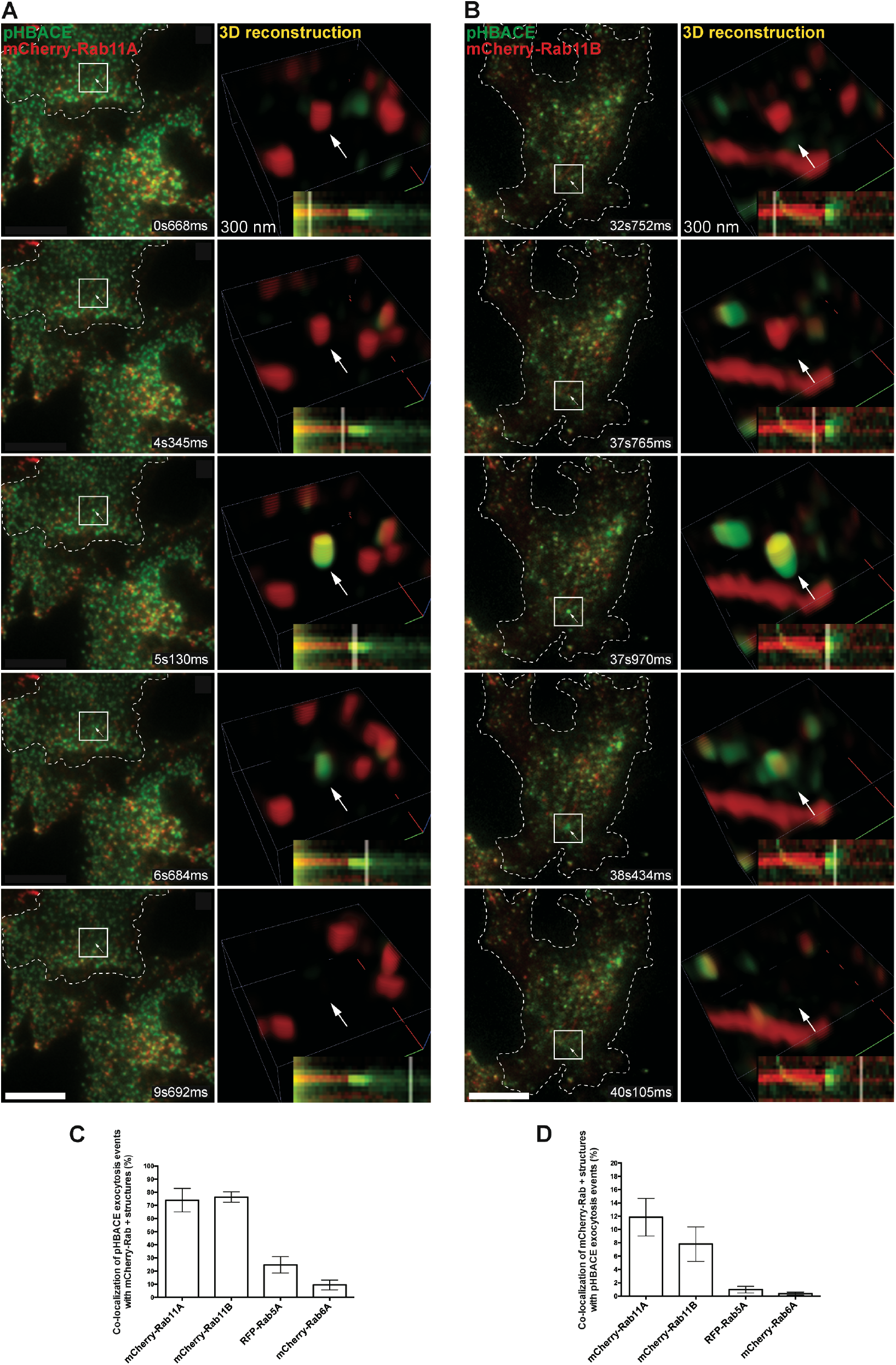
RAB11 participates in exocytosis of BACE1-containing vesicles. HeLa cells co-expressing pHBACE1 and mCherry-RAB11A **(A)** or mCherry-RAB11B **(B)** were imaged using TIRF microscope. Image series corresponding to simultaneous two colors multiangle TIRF image stacks were recorded at one stack of 12 angles every 360 ms (at 30 ms exposure time). 3D reconstructions were performed on the first 300 nm in depth of the cells using a 30-nm axial pixel size. 3D reconstruction of representative exocytotic events are shown (right), including kymographic insets to show time defined events. **(C, D)** Quantification of the co-localization between pHBACE exocytotic events and several mCherry-RABs in cells as described above (mean ± SEM, n= XXX cells). * p= 0.01 (Student’s t-test). Bars: 10 µm

### RAB11 controls BACE1 transport from the sorting endosome

To further investigate the role of RAB11 in BACE1 intracellular trafficking, we depleted RAB11A and RAB11B from HeLa cells using a siRNA duplex designed to target both isoforms (siRAB11A/B, Figure S2B) prior to ectopic expression of BACE1-GFP. Cells were then incubated for 30 minutes with transferrin coupled with Alexa Fluor 647 (Tf-A647) in order to label the early/recycling endosomal pathway. In control cells (siLuciferase), the majority of tubulo-vesicular structures containing BACE1-GFP co-localized with internalized Tf-A647, indicating that they are of endosomal nature (Figure 3A, top row). In siRAB11A/B-treated cells, BACE1-GFP associated with very elongated tubular endosomal structures also positive for internalized Tf-A647 (Figure 3A, bottom row). Similar observation was made upon expression of mCherry-tagged dominant negative mutant of RAB11A or RAB11B (Movies S4 and S5). Interestingly, depletion of RAB11A or RAB11B alone does not induce the appearance of elongated tubular structures contrary to double knock-down (Figures S2A-C), suggesting that the two isoforms work in concert to regulate the morphogenesis of BACE1-containing carriers.

**Figure 3:**
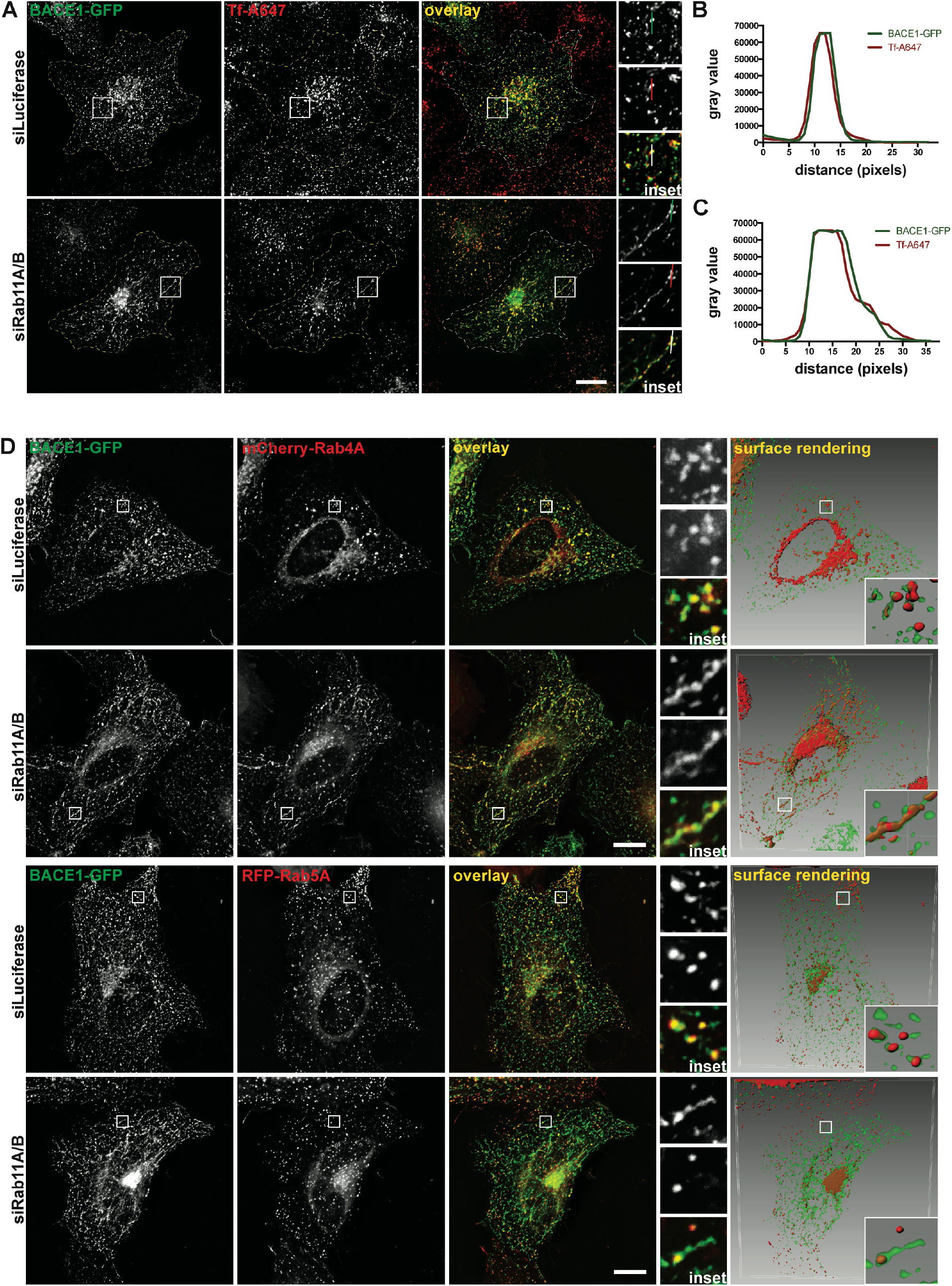
RAB11 controls BACE1 transport at the level of the sorting endosome. **(A)** HeLa cells expressing BACE1-GFP were treated with control or RAB11A/B siRNA for 72h and incubated with Tf-A647 (red). Representative images are displayed. Higher magnifications (boxes) are shown on the right. Line profiles of the BACE1-GFP (green) and Tf-A647 fluorescence intensities (arbitrary units) along the white line. **(B)** HeLa cells co-expressing BACE1-GFP and mCherry-RAB4A and treated with control or RAB11A/B siRNA. Representative images are displayed. Higher magnifications (boxes) are shown on the right. Surface rendering treatment of images (right) highlights the 3D organization of the BACE1-RAB5A positive structures. **(C)** HeLa cells co-expressing BACE1-GFP and RFP-RAB5A and treated for 3 days with control or RAB11A/B siRNA. Representative images are displayed. Higher magnifications (boxes) are shown on the right. Surface rendering treatment of images (right) highlights the 3D organization of the BACE1-RAB4A positive structures.

To better characterize the tubular structures induced by RAB11 depletion, BACE1-GFP was expressed in the presence of RAB5A or RAB4A, two RAB GTPases that associate with EEs/SEs (Sönnichsen et al., 2000);(Stenmark, 2009). As expected, BACE1 transited in control cells through EEs/SEs decorated with both RAB5 and RAB4 (Figure 3D). In siRAB11A/B-treated cells, RAB5 did not associate with the elongated tubular structures containing BACE1-GFP (Figure 3D, bottom row). In contrast, RAB4A was present in these structures.

Altogether, these data suggest that RAB11A and RAB11B regulate the sorting of BACE1 at the level of EEs/SEs. Knowing that RAB4 controls a fast recycling loop from SEs to PM (Sluijs et al., 1992), they also suggest that RAB4 can take over RAB11 recycling function in its absence.

### KIF13A is involved in BACE1 sorting and recycling

We have previously shown that KIF13A cooperates with RAB11 to generate RE tubules from SEs and that KIF13A/B positions RAB11-positive structures to the tips of the cells (Delevoye et al., 2014). To test whether KIF13A is involved in BACE1 recycling, BACE1-GFP and KIF13A-mCherry were co-expressed in HeLa cells. In control cells KIF13A-mCherry was enriched at the cell periphery, consistent with its role in the positioning of RAB11 positive endosomes along microtubules (Delevoye et al., 2009);(Delevoye et al., 2014);(Jani et al., 2022). Interestingly, BACE1-GFP also accumulated at the cell periphery where it co-localized with KIF13A (Figure 4A). Using time-lapse multicolor live-cell imaging, some vesicles co-labelled with BACE1-GFP and KIF13A-mCherry were observed moving together at the cell periphery and in the cell center (Movie S6). Similarly, cells expressing mCherry-KIF13B, a close homolog of KIF13A co-distributing in RE tubules with Rab11A and KIF13A (Jani et al., 2022), displayed peripherally BACE1-GFP-structures positive for KIF13B (Figure 4A). To further validate the role of KIF13 in BACE1 intracellular trafficking, BACE1-GFP was co-expressed with a mCherry-tagged dominant-negative mutant of KIF13A, containing the stalk (S) (residues 351-1306) and tail (T) (1307-1770) domains but lacking the motor domain. As already shown, this mutant exhibits a peculiar pattern of expression and forms abnormally enlarged endosomal structures that did not tubulate (Delevoye et al., 2014). BACE1-GFP co-distributed with ST-KIF13A-positive enlarged structures (Figure 4A, bottom panel), which consist in SEs, as they were decorated with RFP-RAB5A (Figure S3A). Depletion of KIF13A also induced the formation of abnormally enlarged Tf-containing SEs where BACE1-GFP accumulated (Figure 4B and S3B).

**Figure 4:**
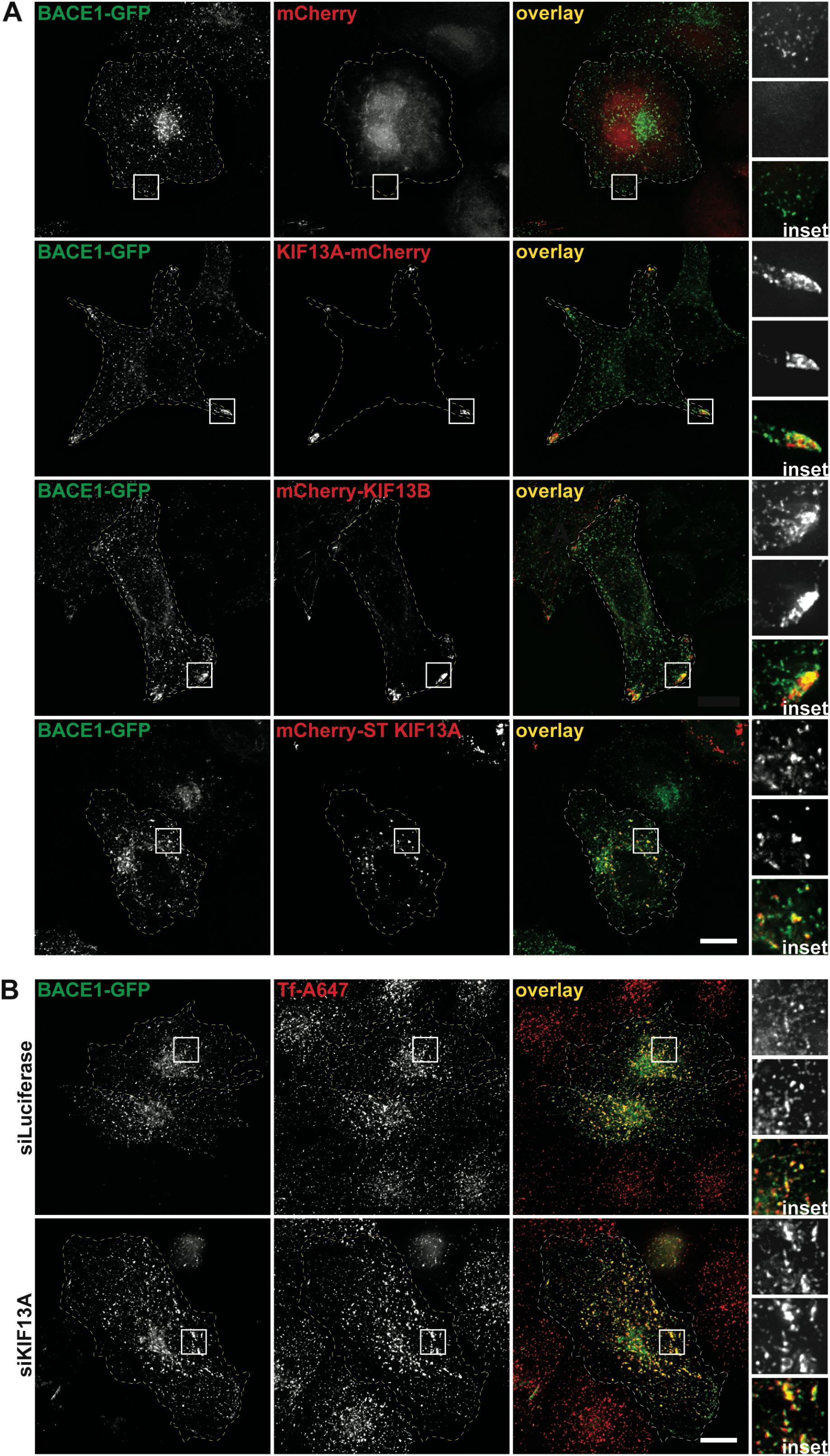
KIF13 contributes to BACE1 endosomal sorting. **(A)** HeLa cells co-expressing BACE1-GFP and mCherry, KIF13A-mCherry, mCherry-KIF13B or mCherry-ST-KIF13A. Representative images are displayed. Higher magnifications (boxes) are shown on the right. **(B)** HeLa cells expressing BACE1-GFP were treated with control or KIF13A siRNA for 3 days and incubated for 30 min with Tf-A647. Representative images are displayed. Higher magnifications (boxes) are shown on the right.

The above data thus support a role of KIF13 in the regulation of BACE1 sorting and recycling. Interestingly, KIF1A, another member of the kinesin-3 family, was shown to mediate axonal transport of BACE1 (Hung and Coleman, 2016). This suggests that the kinesin-3 family of plus end-directed motors may represent bandmasters of BACE1 intracellular trafficking orchestration.

### KIF13-dependent endosome biogenesis machinery is essential for *amyloidogenesis*

SEs function as a trafficking hub from where internalized cargo transport routes diverge. Both APP and BACE1 transit through SEs after internalization from the plasma membrane and many studies described SEs as a major compartment for APP β-cleavage (Toh and Gleeson, 2016). We reasoned that since alteration of KIF13 activity and expression leads to BACE1 accumulation in SEs, this might impact APP processing and Aβ generation. To address this point, we first depleted KIF13A from H4 cells, a neuroblastoma cell line of which APP fragments resulting from its processing can be biochemically monitored (Cavieres et al., 2015) (Figure 5A). Prior to cells lysis, control and KIF13A-depleted cells were treated with γ-secretase inhibitor to increase the amount of the C99 fragment resulting from APP β-cleavage. Cell lysates were assayed by western-blot analysis using an antibody directed against amino acids 4 to 10 of Aβ (Figure 5B). Compared to control cells, KIF13A depleted cells exhibited around 50% increase in C99 intracellular levels (Figure 5B), indicating that β-cleavage is increased when KIF13 expression is downregulated. To further confirm the impact of KIF13 depletion on APP processing and amyloidogenesis, Neuro2a cells were treated with KIF13A or/and KIF13B siRNAs prior to A_1-40_ ELISA assays on cell supernatants. Compared to control cells, the supernatants of Neuro-2a cells depleted of KIF13A, KIF13B or both together exhibited 25 to 60% higher levels of secreted A_1-40_ (Figure 5C). Efficiency of KIF13A/B depletion using specific siRNAs was validated using RT PCR (Figure S4).

**Figure 5:**
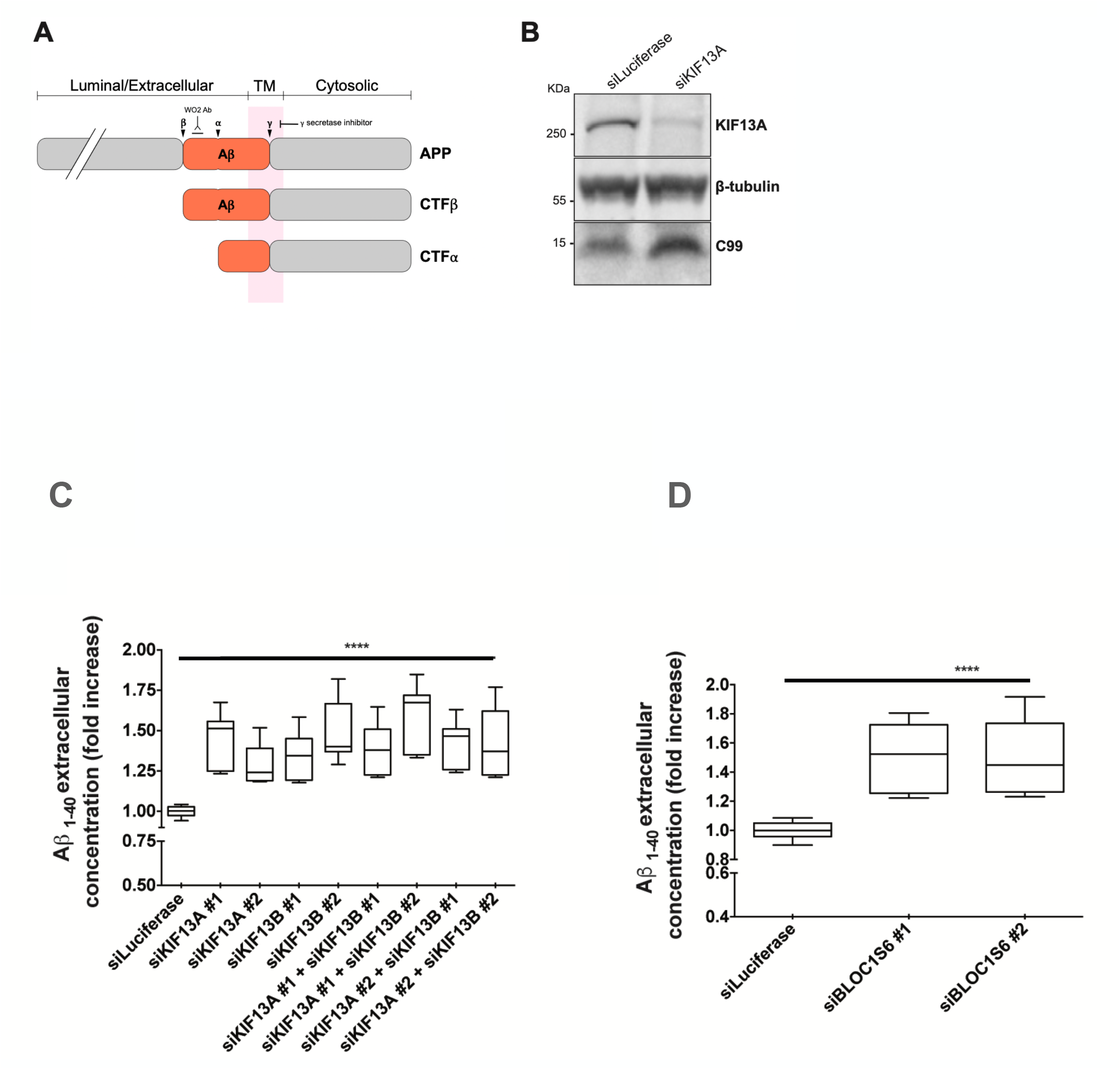
KIF13-dependent endosome biogenesis machinery is essential for amyloidogenesis. **(A)** Schematic of the different domains of APP, CTFb and CTFa. **(B)** H4 cells were treated with control or KIF13A siRNAs. C99 was revealed by Western blot analysis using anti-C99 antibody. **(C)** A_1-40_ ELISA assays on cell supernatants of Neuro2a cells were treated with control or KIF13A and/or KIF13B siRNAs. **(D)** A_1-40_ ELISA assays on cell supernatants of Neuro2a cells were treated with control or BLOC-1 siRNAs. * p= 0.01 (Student’s t-test).

The formation of tubular recycling endosomes from SEs relies on several effectors, including BLOC-1 (Biogenesis of Lysosome-related Organelle Complex 1) that cooperates with KIF13A (Delevoye et al., 2016);(Jani et al., 2022). Similarly to KIF13A/B depletion, BLOC-1 depletion leads to a 50% increase in A_1-40_ secretion (Figure 5D).

In conclusion, our results indicate that the recycling endosome biogenesis machinery, including RAB11, KIF13 and BLOC-1, coordinate BACE1 endosomal sorting to recycling tubules and hence regulate APP β-cleavage and Aβ production. They support the hypothesis that sorting endosomes represent a major location for amyloidogenic processing of APP.

## MATERIAL AND METHODS

### Cells

HeLa, Neuro2A and H4 cells were grown at 37°C with 5% CO2 in Dulbecco’s Modified Eagle Medium (DMEM, high glucose, GlutaMAX, Life Technologies) or DMEM-F12 (Gibco), respectively, supplemented with 10% fetal calf serum (FCS, GE Healthcare and Eurobio), pyruvate sodium 1mM (Life Technologies), penicillin and streptomycin 100 U/mL (Life technologies and Gibco).

### Plasmids

The plasmids encoding the following fusion proteins were used: GFP-/mCherry RAB11A or -RAB11B, KIF13A-GFP, KIF13A-mCherry, mCherry-KIF13B, Lamp1-GFP (C. Delevoye, Institut Curie); mCherry-RAB1A, mCherry-RAB4A, mCherry-RAB5A, BACE1-GFP, pHBACE (B. Goud, Institut Curie, France); APP-mCherry (P. Burgos, Universidad San Sebastian, Chile).

### DNA and RNA transfection

HeLa, Neuro2A and H4 cells were transfected 24 to 48 h before observation with calcium phosphate or with X-tremeGENE9 (Roche), following the manufacturer’s instructions. For RNA interference experiments, cells were transfected with the corresponding siRNA (RAB11A, RAB11B, BLOC-1, KIF13A, KIF13B or Luciferase) using Lipofectamine RNAiMAX (Invitrogen), following the manufacturer’s instructions. The sequences of the siRNAs used in this study: siRNA Luciferase, siRNA against Human RAB11A and Human RAB11B, KIF13A and KIF13B, and BLOC-1 are described in (Delevoye et al., 2014; Delevoye et al., 2016).

### Rat hippocampal neurons culture and transfection

Hippocampal neuronal cultures were performed as described previously (Simon et al., 2013). Briefly, hippocampi of rat embryos were dissected at embryonic day 18, trypsinized and dissociated with a fire-polished Pasteur pipette. Cells were counted and plated on poly-D-lysine-coated glass coverslips and then cultivated in glia-conditioned Neurobasal medium. Neurons were transfected after 7–10 days in vitro (DIV) using Lipofectamine 2000. They were treated and processed at 24h after transfection.

### Immunofluorescence

Cell fixation was performed with paraformaldehyde 3% or 4% (Electron Microscopy Sciences) for 15 min at room temperature. For permeabilization, cells were incubated in PBS supplemented with BSA 2 g/L and saponin 0.5 g/l for 10 min at room temperature. Internalization assays of Tf-A647 were performed as described in (Delevoye et al., 2014). Coverslips were mounted in Mowiol and examined under a 3D deconvolution microscope (Leica DM-RXA2), equipped with a piezo z-drive (Physik Instrument) and a 100x1.4NA-PL-APO objective lens for optical sectioning. 3D or 1D multicolor image stacks were acquired using the Metamorph software (MDS) through a cooled CCD-camera (Photometrics Coolsnap HQ).

### Live cell imaging

For spinning-disk confocal time-lapse imaging, the samples were imaged at 37°C in a thermostat-controlled chamber (Life Imaging Service) using an inverted Eclipse Ti-E microscope (Nikon) equipped with a spinning disk X1confocal head (Yokogawa), a 100x NA1.4 oil objective, and either an EMCCD Ultra897 iXon camera (Andor Technology) or a CCD CoolSnapHQ2 camera (Photometrics). The imaging was controlled by Metamorph 7.8 (Molecular Devices).

For TIRF microscopy, the samples were imaged using Leibovitz’s medium (Life Technologies) at 37°C in a thermostat-controlled chamber. Eclipse Ti inverted microscopes (Nikon) equipped with either a fixed angle TIRF module (Nikon) or an ILAS2 azimuthal TIRF module (GATACA systems), a 100x TIRF objective (NA=1.49), a beam splitter (Roper Scientific) and an Evolve 512 EMCCD camera (Photometrics) were used. In the latter case, MA-TIRFM was performed (10 to 12 angles). Restoration and visualization were based on modeling PSF at the various angles, as previously described (Boulanger et al, 2014) All acquisitions were driven by Metamorph (Molecular Devices).

### Cell lysis, SDS-PAGE, and Western-blotting

Cells were washed three times in ice-cold PBS, scraped and then lysed in a buffer containing 150 mM NaCl, 50 mM Tris-HCl pH 7.5, 1% Nonidet-P40 (Sigma). Protein concentrations were determined by Quick Start™ Bradford 1x Dye Reagent (Bio-Rad). Equal amounts of proteins were reduced with 1x loading buffer containing 6% β-mercaptoethanol and resolved on 10% SDS-PAGE. Proteins were transferred onto nitrocellulose Protran BA 83 membrane (Life science), processing for immunoblotting. HRP-conjugated secondary antibodies associated signal was detected with ECL system and an enhanced chemiluminescence system (ChemiDoc Touch System, Bio-Rad). Quantification of the corresponding signal obtained was done using Image lab software.

### Image analysis and quantification

In Figure 2C, 2D, colocalization between fixed vesicles in two different channels was determined manually using Image J software (Synchronize windows). In Figure S2C, the % of cells containing BACE1-positive elongated structures was determined manually using Image J software. In Figure S1F, colocalization was measured with Pearson’s coefficient using Image J software.

### Statistical analysis

All data were generated from cells pooled from at least 3 independent experiments represented as (n), unless mentioned, in corresponding legends. Statistical data were presented as means ± standard error of the mean (S.E.M.). Statistical significance was determined by Student’s t-test for two or three sets of data using Excel, no sample was excluded. Cells were randomly selected. Only P-value <0.05 was considered as statistically significant.

## ACKNOWLEDGEMENTS

The authors gratefully acknowledge the Cell and Tissue Imaging Facility (PICT-IBiSA), Institut Curie, a member of the French National Research Infrastructure, France-BioImaging (ANR-10-INBS-04-01). The authors also thank the recombinant antibody platform of the Institut Curie. The work performed in B.G. laboratory was supported by a grant from France Alzheimer association and an ERC (European Research Council) advanced grant (project 339847 ‘MYODYN’). The Goud and Serpico/STED team are members of the Labex Cell’n’Scale (11-LBX-0038) and Idex Paris Sciences et Lettres (ANR-10-IDEX-0001-02 PSL). Serpico/STED is a member team of the French National Research Infrastructure, France-BioImaging (ANR-10-INBS-04-07).

## Authors Contributions

JB.B. carried out all the experiments; Z.L. performed the TIRF experiments and corresponding analysis presented in Figure 2; D.L. prepared the samples for experiments presented in Figure. 1B. V.F. performed the 3D analysis presented in Figure 3B. JB.B., Z.L., S.B., D.L., A.T., Z.L., J.S., C.D., B.G., S.M. designed and interpreted the experiments. JB.B., C.D., B.G. wrote the manuscript. JB.B., Z.L., S.B., D.L., A.T., Z.L., J.S., C.D., B.G., S.M. edited the manuscript. B.G and S.M. secured funding.

## Figure Legends

**Supplementary figure S1**

Representative Images of HeLa cells co-expressing either **(A)** GFP-RAB11A and mCherry-RAB11A, BACE1-GFP and mCherry-RAB1A, **(C)** APP-mCherry and GFP-RAB11A, APP-mCherry and Lamp1-GFP. (A-D) Higher magnifications (boxes) are shown on the right. **(C)** Line profiles of the APP-mCherry (red) and Lamp1-GFP fluorescence intensities (arbitrary units) along the line. **(D)** Colocalization (Pearson’s coefficient) between GFP-RAB11A and mCherry-RAB11A, BACE1-GFP and mCherry-RAB11A, BACE1-GFP and mCherry-RAB11B, BACE1-GFP and mCherry-RAB1A, APP-mCherry and GFP-RAB11A (mean ± SEM,).

**Supplementary figure S2**

**(A)** HeLa cells expressing BACE1-GFP were either non treated or treated for 3 days with siRNAs control sequence or siRNAs targeting RAB11A, RAB11B, RAB11A+RAB11B, RAB11A/B. **(B)** HeLa cells were treated as indicated in (A) and processed for western-blotting. RAB11 signal was revealed using an anti-RAB11 antibody. Beta-tubulin signal was used as a loading control. **(C)** Quantification of the % of cells containing BACE1-positive tubules in cells treated as indicated in (A).

**Supplementary figure S3**

**(A)** Representative Images of HeLa cells co-expressing GFP-ST-KIF13A and RFP-RAB5A. Higher magnifications (boxes) are shown on the right. **(B)** Representative Images of HeLa cells treated 3 days with control or KIF13A siRNA and then co-transfected with BACE1-GFP and RFP-RAB5A. Higher magnifications (boxes) are shown on the right.

**Supplementary figure S4**

HeLa cells were treated with either control, KIF13A, KIF13B siRNAs for 3 days. The level of KIF13A/B expression was measured using RT-PCR.

**Supplemental movies:**

S1: BACE1-GFP + mCherry-RAB11A in rat hippocampal neurons

S2: 3D TIRF + plot profile + kymograph + event number (see Boulanger et al., PNAS, 2015) for pHBACE + mCherry-RAB11A

S3: 3D TIRF + plot profile + kymograph + event number (see Boulanger et al., PNAS, 2015) for pHBACE + mCherry-RAB11B

S4: BACE1-GFP + mCherry RAB11A S5: BACE1-GFP + mCherry-RAB11B

S6: BACE1-GFP and KIF13A-mCherry

