## Supplementary figures and images for "The recycling endosome biogenesis machinery coordinates BACE1 endosomal sorting and amyloid-β production"

### Supplemental Figure 1

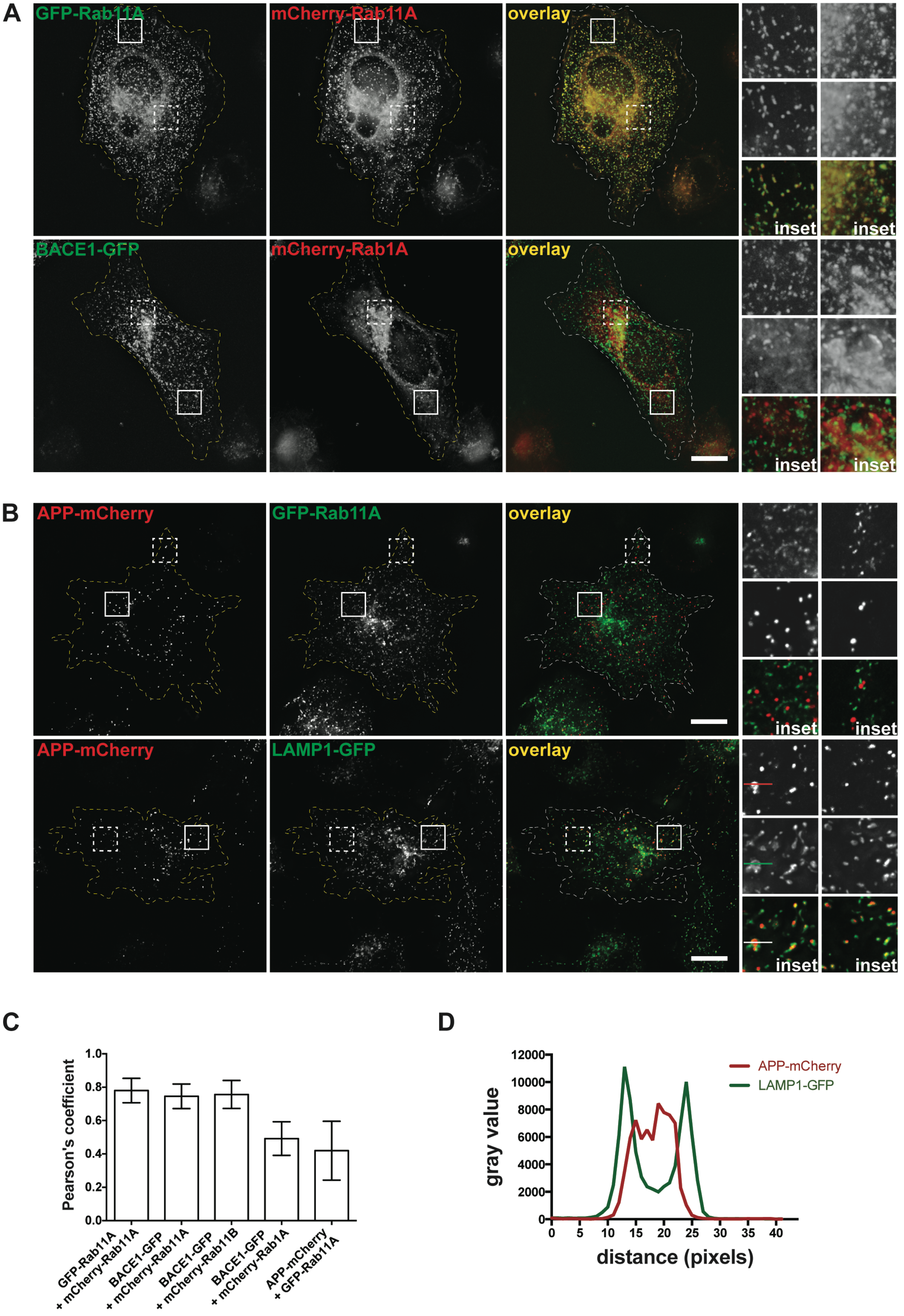

### Supplemental Figure 2

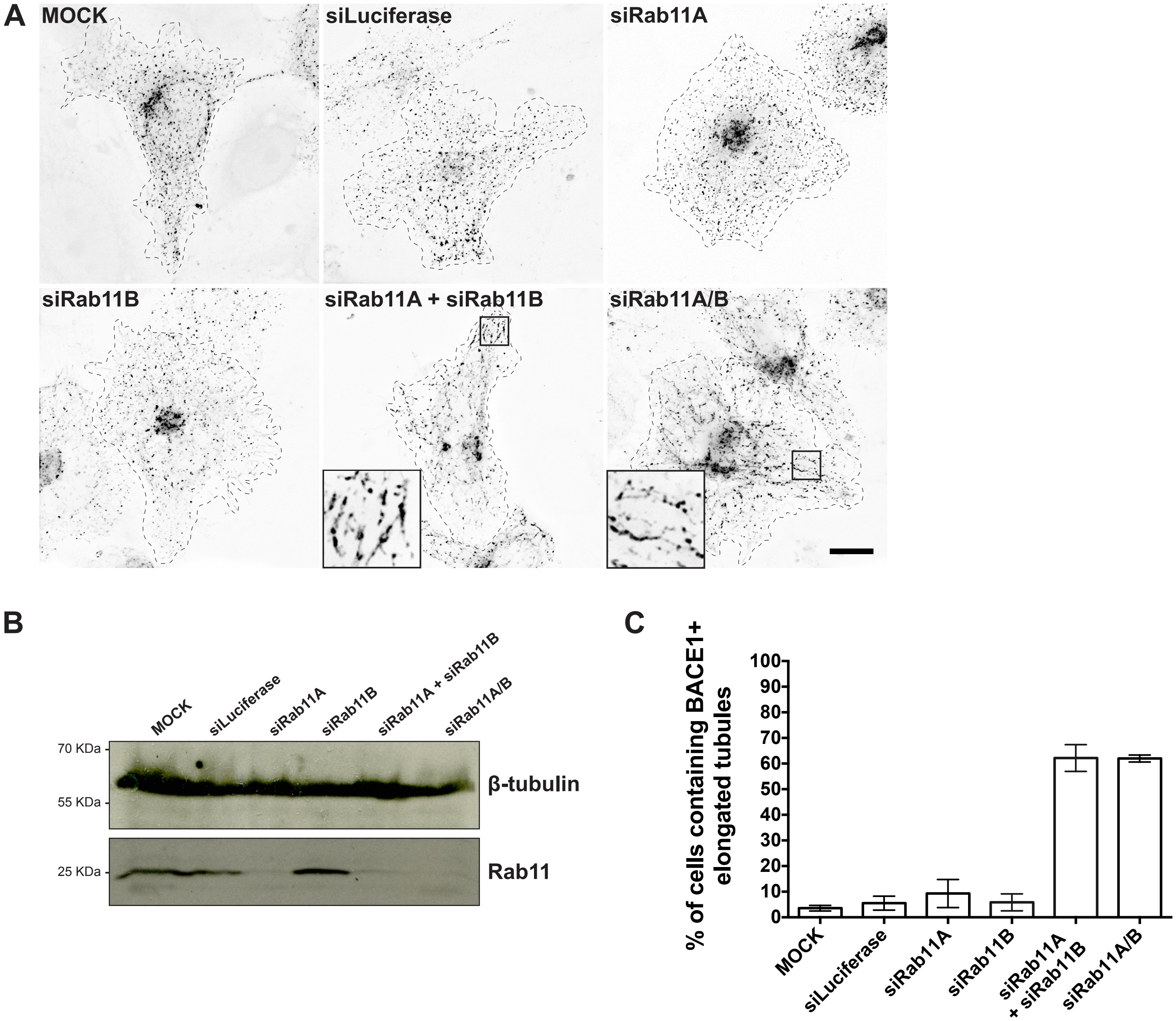

### Supplemental Figure 3

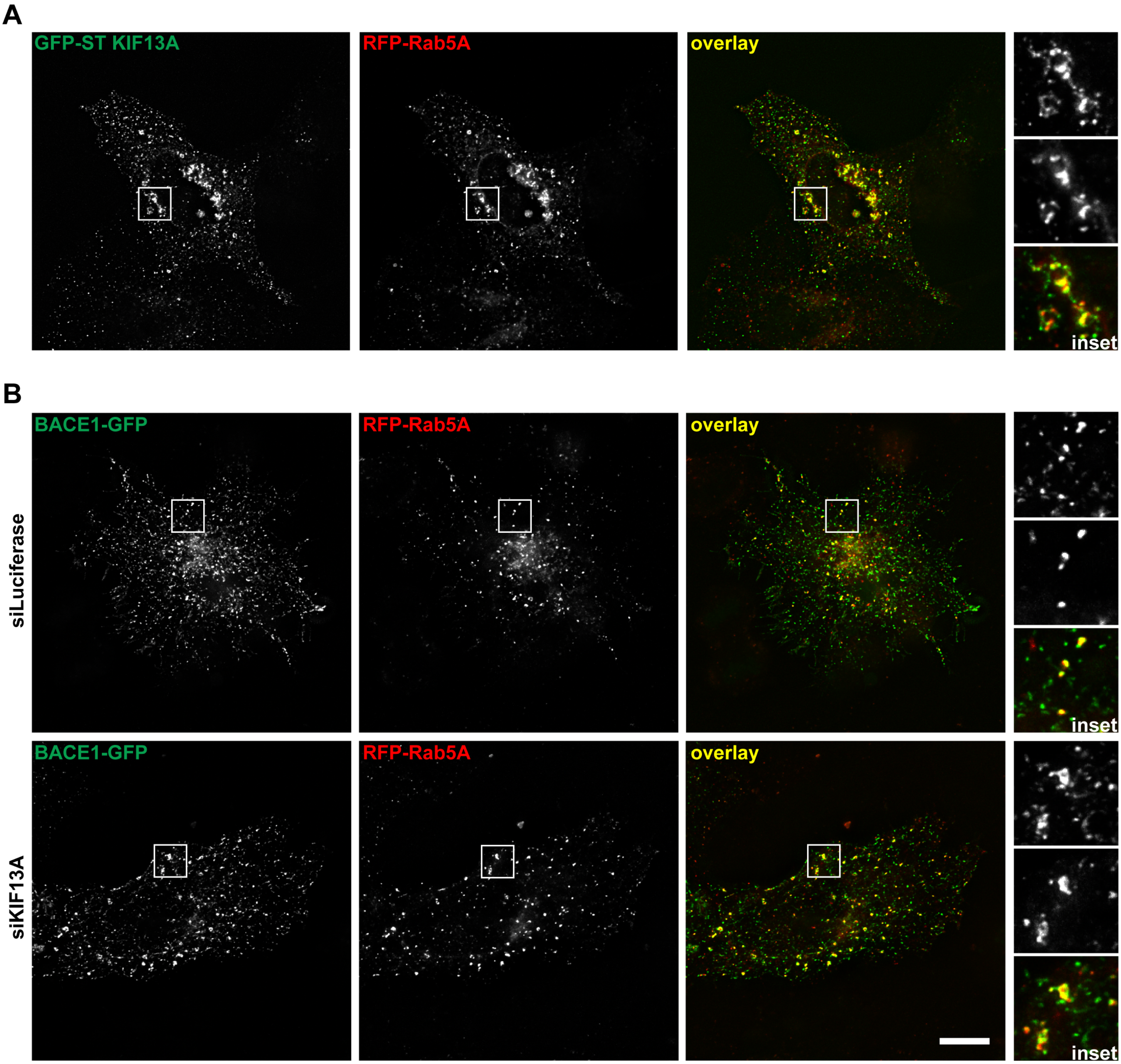

### Supplemental Figure 4

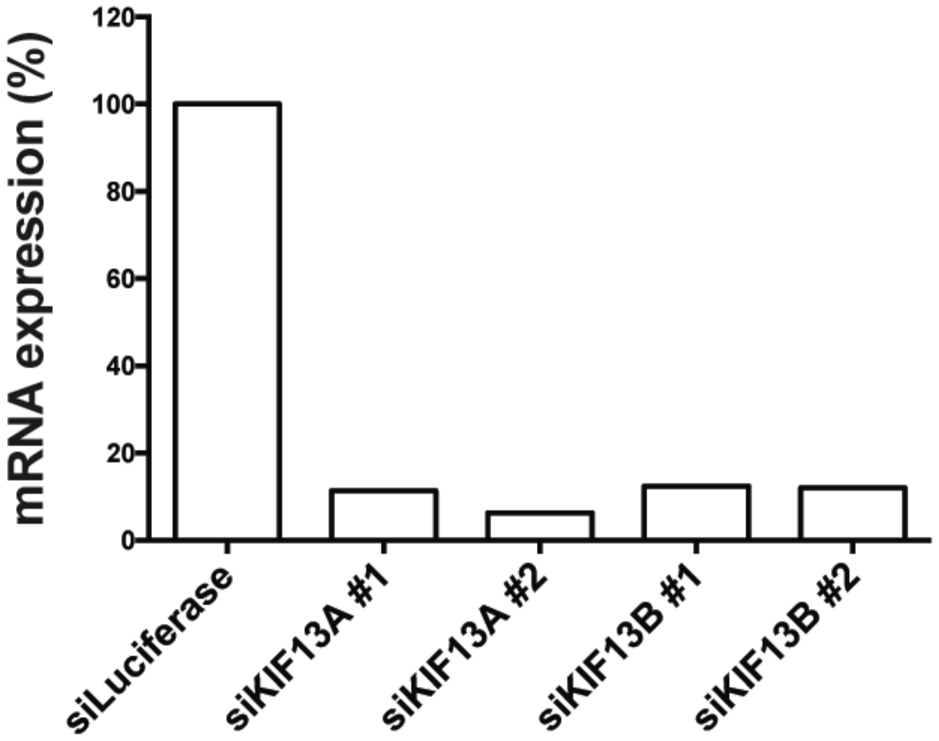
